# Tying up loose ends: Recovering thousands of missing telomeres from Streptomyces and other Streptomycetaceae genomes

**DOI:** 10.1101/2025.10.14.682034

**Authors:** David Faurdal, T.J. Booth, Tilmann Weber, Tue Sparholt Jørgensen

## Abstract

Members of the Gram-positive *Streptomycetaceae* family of bacteria have linear chromosomes and carry linear plasmids, which end in telomeres bound by proteins. In a large-scale analysis of 762 linear complete genomes, we discovered that the telomeres were truncated in most assemblies, as they are not captured by Oxford Nanopore sequencing. To address this issue, we present Telomore, a tool to reconstitute this missing telomeric sequence using ONT and Illumina data. In the studied dataset, Telomore increased detection of archetypal telomeres from 0% to 37%, which could be near the occurrence rate in nature. Combining these reconstituted telomeres with previously published telomeres and all complete *Streptomycetaceae* RefSeq genomes, we created a compendium of more than 2000 telomeres. Similarity-based clustering identified 137 telomere clusters. We find that 78% of Telomore-extended chromosomes encode both telomeres, while this is only the case for 15% of comparable RefSeq chromosomes. Therefore, most assignments of “complete” to *Streptomycetaceae* are erroneous. Finally, we mined the 762 genomes for known telomeric maintenance proteins and used those to identify a plasmid-specific archetypal telomere and to identify a previously unidentified protein family likely involved with the maintenance of Sg2247-class telomeres. Together, these results highlight a common issue assembling complete linear Streptomycetaceae genomes and provide a programmatic solution and identify a candidate for a new telomeric protein.

**VISUAL ABSTRACT:** 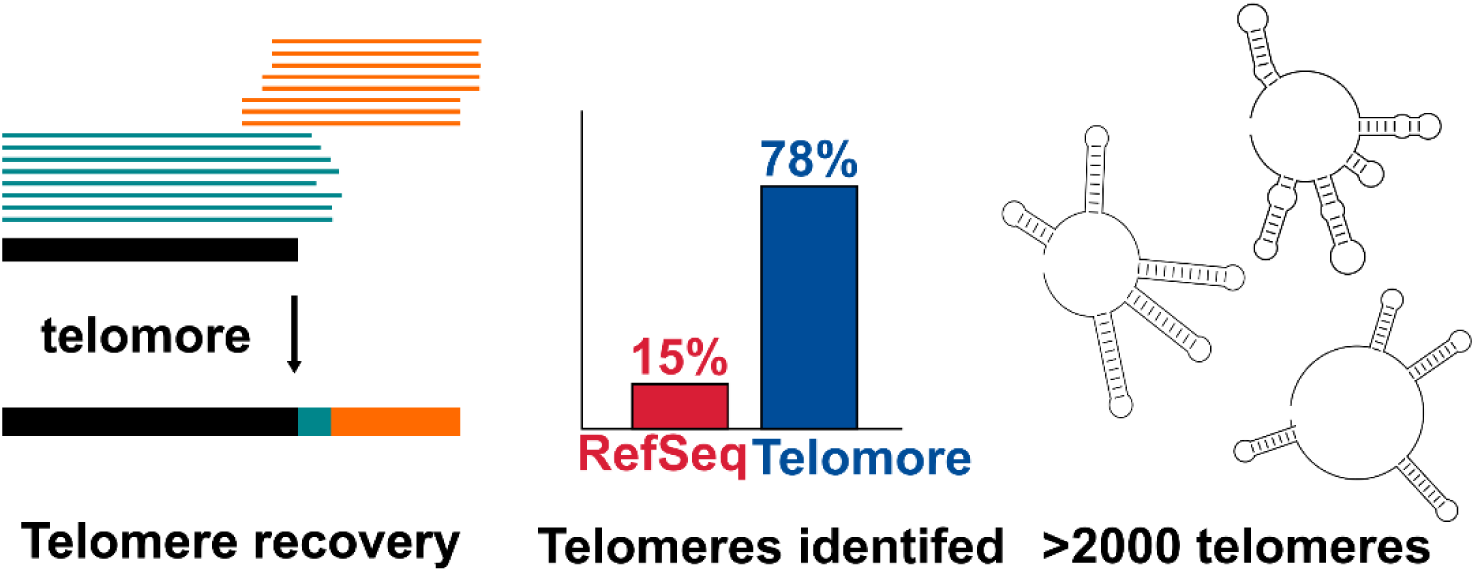

## INTRODUCTION

Species in the bacterial family *Streptomycetaceae* have linear chromosomes and often carry linear plasmids. To maintain a linear replicon an organism needs to circumvent the end-replication problem, where the 5’-end of the lagging strand is not fully replicated, leaving an overhanging 3’-end. In *Streptomycetaceae*, the overhanging 3’-strand folds into a secondary structure, which is then bound and repaired by telomeric proteins. This is a unique approach to maintaining linear replicons and understanding the diversity of these telomeres can provide practical benefits for i) the evaluation of genome assembly completeness, as each end of a linear replicon should contain a telomere. ii) the development of synthetic biology tools based on linear replicons.

The ‘archetypal’ telomere system in *Streptomycetaceae*, primarily studied in the genus *Streptomyces*, is the most well characterized and is also thought to be the most widespread. Here, the overhanging telomere is bound by telomere-associated protein (Tap), which in turn recruits terminal protein (Tpg) to the terminus, where they perform end-patching (1–3). Archetypal telomeres have a conserved sequence, which contains several palindromic sequences predicted to fold into a single-stranded secondary structure during end-patching (4). The first three palindromes form loops of 5’-GGA-3’, 5’-TGC-3’ and 5’-TGC-3’ and are highly conserved while the subsequent palindromes show decreasing conservation. These conserved palindromes serve as the binding target of Tap/Tpg. Specifically, Palindromes II and III are bound by Tap (1), following which Tpg is recruited to the terminus of Palindrome I. Together, the Tap-Tpg complex primes the synthesis of a 13 nt primer (the reverse complement of palindrome I) at the 5’-end (2, 3). During this process Tpg is directly deoxynucleotidylated by Tap, which leaves it covalently bound to the 5’-end of the replicon (3) providing resistance to exonuclease degradation. After primer synthesis, one of the housekeeping DNA polymerases, DinB1 or DinB2, uses the 13 nt primer to synthesize the remaining lagging strand (3, 5, 6).

In addition to the archetypal telomere, several non-archetypal telomeres have been described in the literature. In 2021, Algora-Gallardo et al. (7) categorized 108 telomere sequences into six classes of telomeres: the Sco class, composed of archetypal telomeres similar to the chromosomal ends of *Streptomyces coelicolor*; the Scp1 class, composed of telomeres similar to the giant linear plasmid SCP1; the Sg13350 and Sg2247 classes consisting of sequences similar to those found in *Streptomyces griseus* IFO13350 and *S. griseus* 2247; and the pRL1 and pRL2 classes, which are each represented by a single member from the telomeres of the two eponymous plasmids. Key features of each class are outlined in Table 1, while the folding of a representative of each class is visualized in Figure 1.

**Figure 1:**
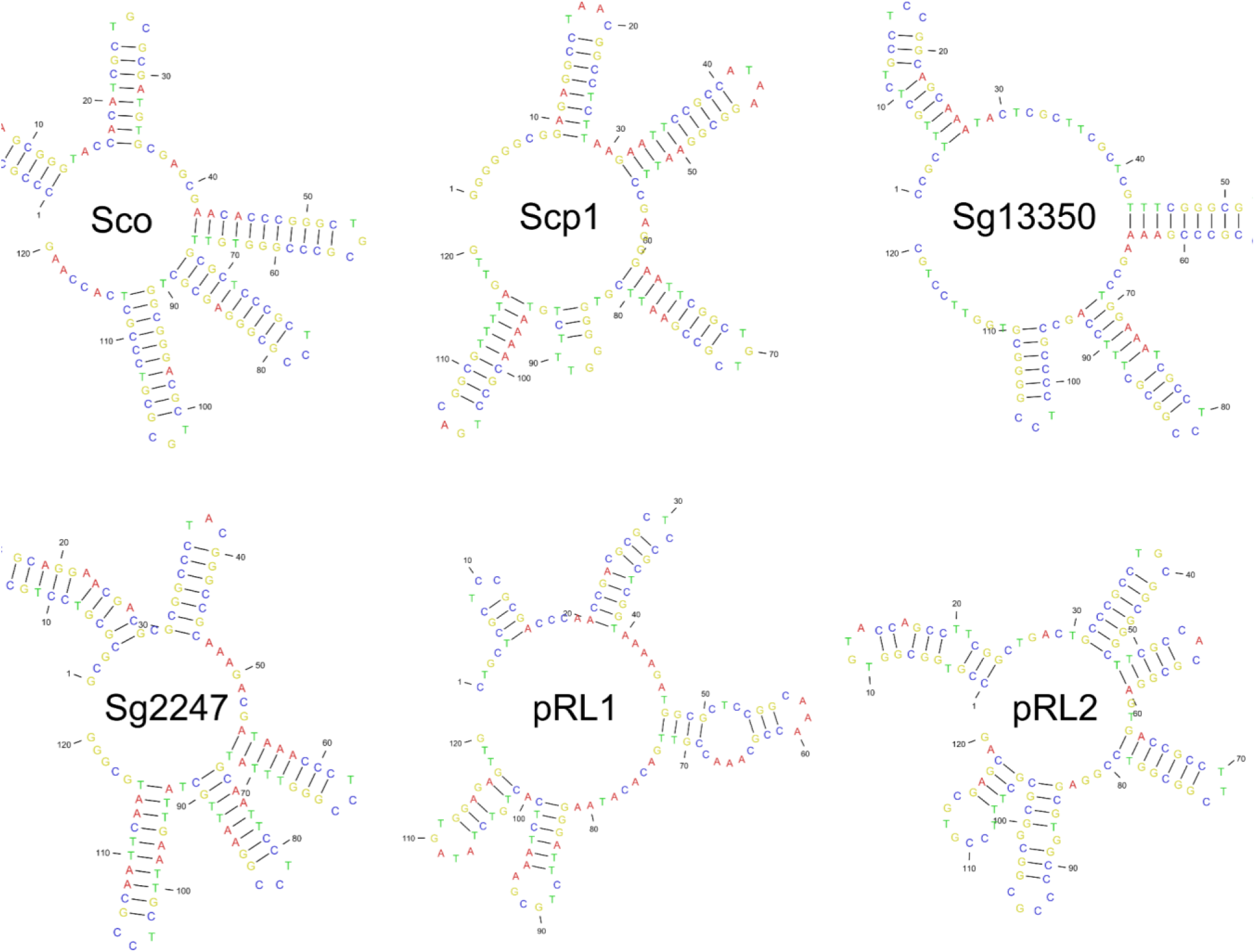
The six classes of *Streptomyces* telomeres. The six classes of telomeres as described by Algora-Gallardo et al. (7)

**Table 1:** Information on the six classes of *Streptomyces* telomeres.

| **Telomere Class** | **Named After** | **Key Features** | **Maintenance proteins** | **References** |
| --- | --- | --- | --- | --- |
| Sco<br>(Archetypal) | *S. coelicolor*,<br>chromosomal ends | Three C's at the end;<br>typical palindrome<br>structure | Tap/Tpg | Algora-Gallardo et al.,<br>2021 |
| SCP1 | Giant linear plasmid<br>SCP1 | G-rich end instead of<br>three C's; palindromes<br>with 4-base loops | Tac/Tpc | Bentley et al., 2004;<br>Huang et al., 2007 |
| Sg13350 | *S. griseus* IFO13350 | Palindromes with 5'-<br>TCC-3' loops | GtpA/GtpB | Ohnishi et al., 2008;<br>Suzuki et al., 2008 |
| Sg2247 | *S. griseus* 2247 | Ends with a GC-repeat. 5'-TCC-3';<br>Palindromes II & III:<br>loop 5'-TAC-3' | TmpA (candidate<br>from this study) | Goshi et al., 2002 &<br>Algora-Gallardo et al.,<br>2021; This study |
| pRL1 | Linear plasmid pRL1<br>from *Streptomyces*<br>*sp.* 44030 | Only known member<br>of its class | Unknown | Zhang et al., 2006 |
| pRL2 | Linear plasmid pRL2<br>from *Streptomyces*<br>*sp.* 44414 | Only known member<br>of its class | Unknown | Zhang et al., 2006 |

**Table 2:** Overview of replicon ends extended by Telomore. . Both: Extended first with Oxford nanopore technologies (ONT) reads and subsequently Illumina reads. ONT only: Extended with ONT reads, but failed extension with Illumina reads. Illumina only: Not extended by ONT reads but extended by Illumina reads. None: Both types of data failed to extend these ends.

| **Replicon Type\extended by** | **Both ONT and Illumina** | **ONT only** | **Illumina only** | **None** |
| --- | --- | --- | --- | --- |
| Chromosome | 1050 | 132 | 236 | 106 |
| Plasmid | 619 | 52 | 197 | 78 |
| Unclassified Extrachromosomal | 83 | 9 | 49 | 7 |
| Total | 1752 | 193 | 482 | 191 |

**Table 3:**
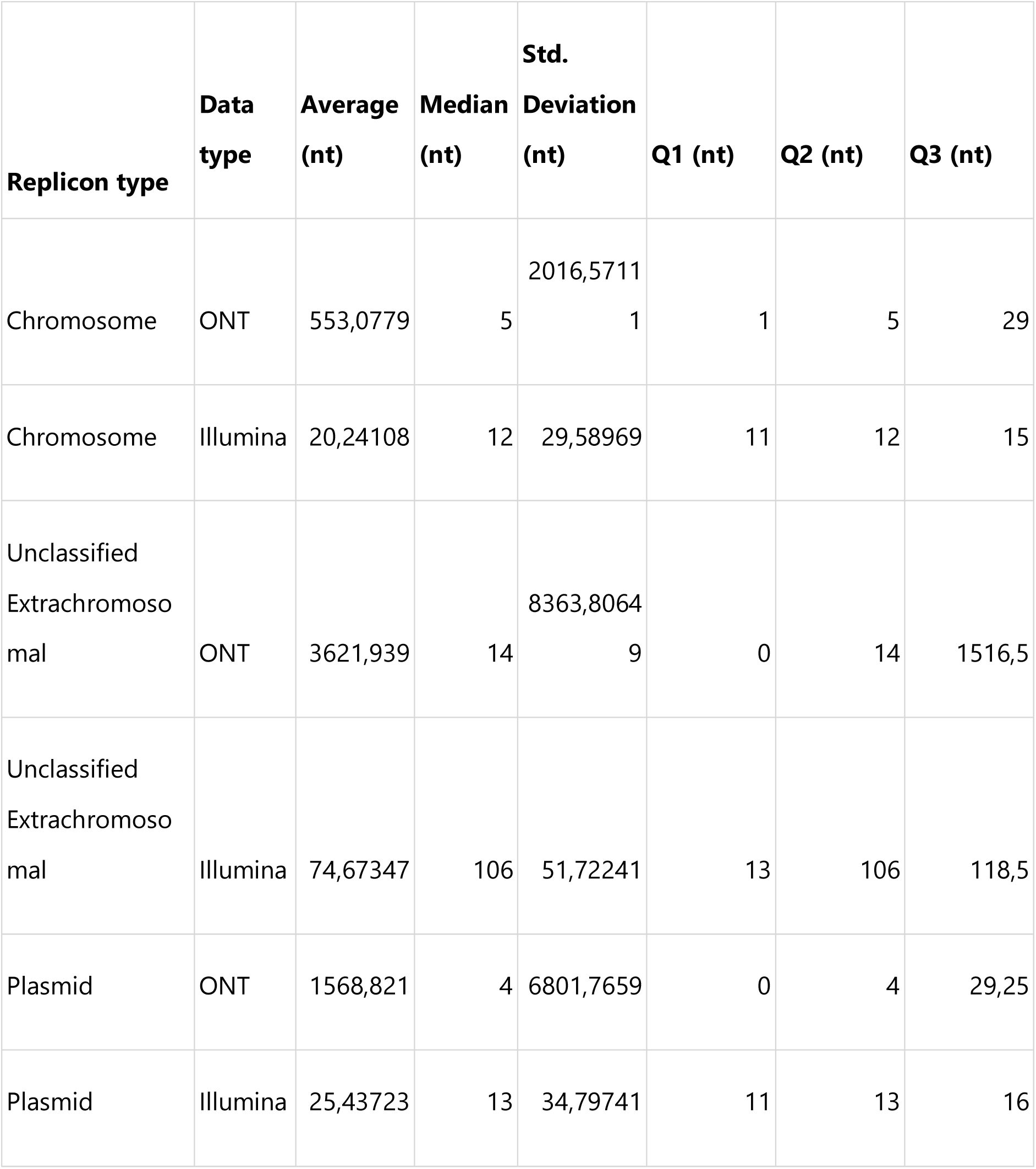
Summary statistics for extension of replicons by Telomore. . Replicons were first extended with Oxford Nanopore Technologies (ONT) reads. These extended contigs were then extended with Illumina reads.

At the time of Algora-Gallardo et al.’s original classification, most complete genomes deposited at NCBI did not have an identifiable telomere at each end , which could be due to unknown telomere classes or genome incompleteness (7). In recent years, “long-read first” hybrid assembly has become the norm for accurate and complete assembly of bacterial genomes (8).This has led to a large increase in the available number of complete *Streptomycetaceae* genomes (9). With this in mind, we set out to analyze the telomeres of publicly available genomes.

To explore telomere diversity, we started by analyzing the G1034 collection (9), which consists of 762 linear complete *Streptomycetaceae* genomes, assembled by long-read first hybrid-assembly using Oxford Nanopore Sequencing (ONT) and Illumina sequencing. During inspection of the replicon ends we noticed that an archetypal telomere was often present in the assembly but truncated. This prompted us to question whether or not telomeres are accurately captured by ONT and whether otherwise complete genome assemblies fail to capture the very ends of the replicons.

## MATERIALS & METHODS

### Telomore: A python workflow for extending linear contigs

A python workflow, Telomore [github: https://github.com/dalofa/telomore], was developed to reconstitute the missing telomeric sequence of an otherwise complete assembly using ONT data and Illumina data. Telomore consists of five analysis steps (Figure 2), briefly: i) mapping reads against all contigs in a reference using either minimap2 (10) or Bowtie2 (11). ii) Extending reads (i.e. those that overhang the ends of linear contigs) are extracted using SAMtools (12) iii) the extending reads from each end are used to construct a consensus sequence using either lamassemble(13) or mafft(14) and EMBOSS(15)]; iv) The consensus for each end is aligned to the reference and used to extend it v) all terminally mapped reads are mapped to the new extended reference and used to trim away spurious sequence, based on read-support and/or base quality.

**Figure 2:**
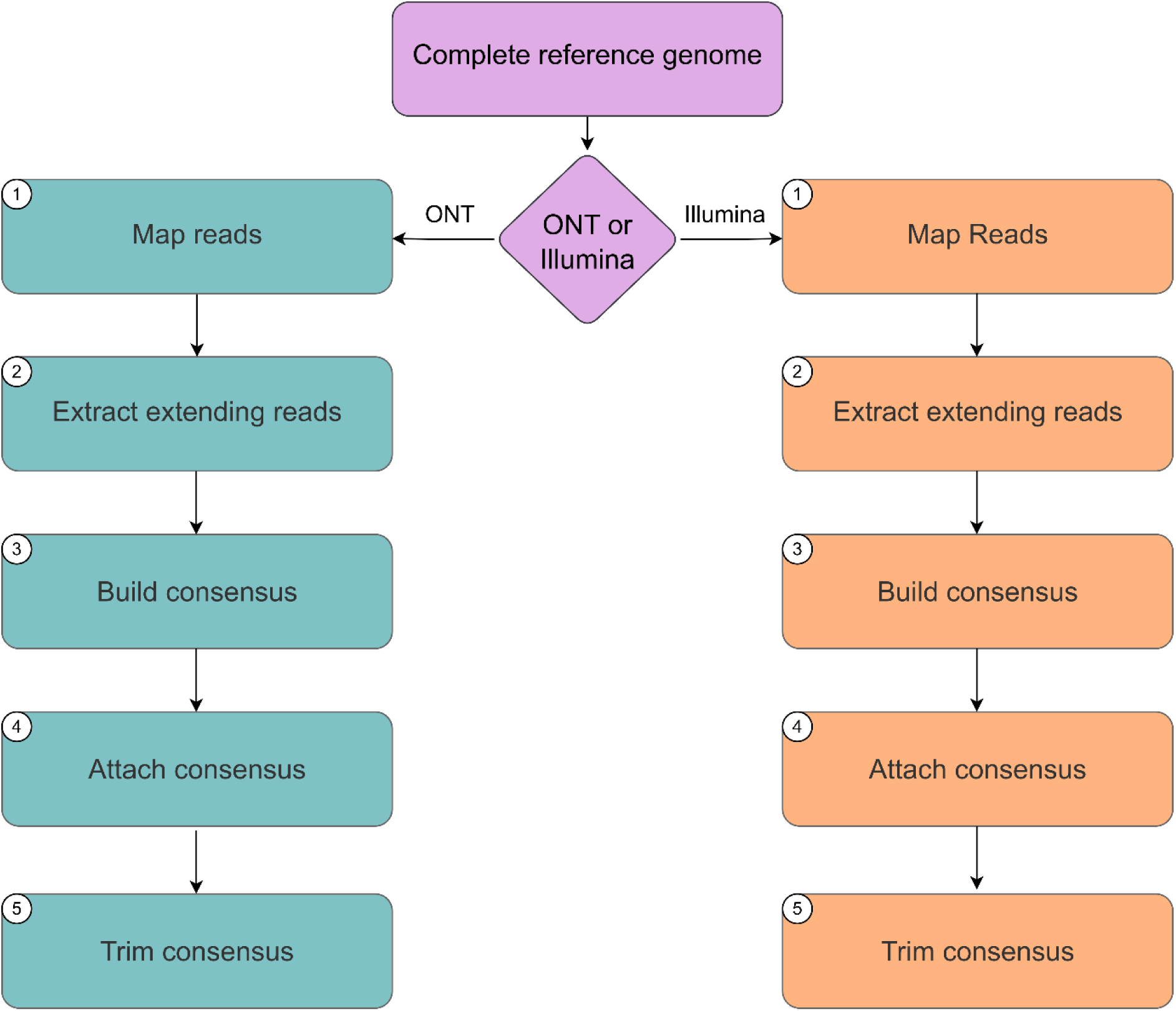
Telomore workflow, see text for description of each step. Steps using Illumina reads are colored orange, while steps using ONT reads are colored teal.

### Reconstituting telomeric sequence using ONT and Illumina data

#### Preparation of Illumina reads

Illumina reads were trimmed using trimgalore (v0.6.5 running cutadapt v2.10) with a minimum length of 100 nt and a minimum quality of Q20(16, 17)(17).

#### Extension by Telomore

The linear replicons from 762 strains, of which 3 chromosomes were excluded due to technical sequence, were subject to extension with Telomore V0.3 (commit: #df04cfd). First linear replicons were extended ONT data. All linear contigs were then extended using Illumina reads and finally these extended contigs were polished using polypolish (18)

#### Detection of archetypal palindrome I

The first or last 90 nt of each replicon were subjugated to BLAST (19) searches using palindrome I (CCCGC**GGA**GCGGG) as query applying `-task blastn-short.` The outputs were then filtered for sequences containing at least 10 of the 13 bases starting within the first 3 bases.

#### Clustering of telomeric sequences

##### Telomere dataset creation and filtering

Only linear replicons are expected to carry telomeres, and each linear replicon has two ends, each carrying a telomere, which can be identical to or different from the other end of the replicon.

We clustered replicon ends from 2665 replicons from 1454 genomes, of which 1123 replicons were downloaded (2025-01-17) from NCBIs Refseqs complete and chromosome level *Streptomycetaceae* genomes and 1532 replicons (1309 of which are linear) were from our own previous work, completed using Telomore. In addition, we included telomeres deposited individually (see supplementary Table S1). The Telomore-extended genome of NZ_CP080029 was included as well as the original sequence.

A small number of replicon ends were found to contain many N’s, these were removed before clustering. Similarly, a number of sequences were found to contain low complexity sequences (e.g. CACACACA or GGGGGGGG for stretches of 20+ nt). The low complexity sequence was primarily but not exclusively found in data previously submitted by our group. Since the low complexity sequences were not expected or observed to cluster, they were left in the analysis.

A single genome was found to consist of 150 contigs: these sequences were removed from analysis and NCBI informed that the genome assembly was not in fact complete.

17 replicon ends were found to contain ONT nanopore technical sequence, namely, rapid barcodes, adapters and barcode flanking sequences. These were removed prior to further analysis and reported to NCBI for removal. Manual inspection showed that in all 17 cases, the technical sequence issue existed before Telomore completion, and so in no case did Telomore create a technical sequence issue.

In total, the dataset used for clustering consists of 5324 sequences, of which 4606 are linear replicon ends or reported telomeres. Tables with an overview of the strains used for clustering and the fasta files used to cluster telomeres can be found at https://github.com/dalofa/telomore_paper/tree/main/clustering. See Supplementary Table S1 for an overview of filtered replicon ends.

### Telomere clustering

The subseq command of Seqkit (v2.9.0) (20) was used to extract the first and last 90nt of each replicon end and the Seqret -srev command (15) used to orient the sequences. For clustering, Vsearch (v2.28.1) (21) was used with the following parameters: --id 0.72 and --iddef 0 to cluster on 72% identity using the CD-HIT clustering algorithm. We then removed clusters with fewer than four members.

### Telomere folding and alignment

To annotate and visualize the folding of the single stranded telomere sequences, we used the CLC Main Workbench (v.22.0.2) Predict Secondary Structure function with a maximum distance between pairs of 25 nt. Visualizations of structure annotation and foldings were also made in CLCMain Workbench. To visualize the within-telocluster variation and class to cluster differences, we aligned the first 120nt of all sequences in each telocluster using the CLC Genomics Workbench (v.12.0.3) batch Create Alignment function.

### Mining for telomere maintenance machinery in the G1034 collection

The Tap (accession: CAC22741.1) and Tpg (accession: CAC22742.1) proteins from *S. coelicolo*r A3(2) were used as queries in BLASTp-searches against all annotated CDSs in the 752 complete *Streptomycetaceae* genomes of the G1034 collection pre-Telomore, as these are fully annotated with publicly available locus tags. Hits with an e-value above 1e-10 or a query coverage below 40% were excluded.

Similarly, Tac (accession: CAC36646.1) and Tpc (accession: CAC36648.1) from the plasmid SCP1 and GtpA (accession: BAG16926) and GtpB (accession: BAG16927) from *S. griseus* IFO1335 were also used as queries to search against the G1034 collection, applying the same thresholds.

### Subset of co-localized gene-pairs

Tap-Tpgs, Tac-Tpcs and GtpA-GtpB were filtered to only include hits where the two genes were co-localized (a 300 nt intergenic distance or smaller) and assigned to the telomere end closest to the gene-pair under the assumption that they will generally tend to co-locate with their cognate telomere. The translated protein sequences were then clustered into families at 70% similarity using CD-HIT(22). Co-occurrence of the proteins with its partner and teloclusters were visualized. To avoid spurious correlations, protein-to-teloclusters supported by less than 2 replicon-ends were filtered from the dataset after manual inspection.

### Identification of protein families associated with an unknown system

Protein coding sequences from 762 genomes were clustered using CD-Hit (22) using a global sequence identity of 0.7. Representatives of the resulting protein “families” were searched against the 762 genomes using blastp, as implemented in DIAMOND(23). This was used to calculate relative frequencies of each protein family in genomes with a known maintenance system (583 genomes) and genomes with an unknown maintenance system (179 genomes). Plotting these results on an X-Y scatter plot enabled the identification of outlying clusters that were overrepresented in systems with an unknown system.

### Data visualization and statistical methods

All scripts used for protein mining and plotting can be found in the github repository https://github.com/dalofa/telomore_paper. All plots were generated using R. The associations between categorical variables (telocluster and protein family) were tested using Cramer’s V using the rcompanion package.

Telomeres are depicted as the 5’-end of assemblies, which in principle is the sequence complementary to what is overhanging during end-patching, as the 5’-end is recessed. We chose this representation as it follows the convention of DNA sequence representation of 5’-3’ direction, reading from left to right.

### Domain detection using InterProScan

Protein sequences were searched for known domains using the InterProScan (24) webserver with default settings.

### Structural modelling of proteins

Protein structures were modeled using Colabfold (25) v1.5.5, which applies Alphafold2. The best ranked model for each protein was compared using the TM-align server(26) with default settings.

## RESULTS AND DISCUSSION

### Missing telomeric sequence can be reconstituted using Telomore

Initial inspection of the termini of linear genomes from the G1034 collection revealed that many chromosomes contained a truncated archetypal telomere. In fact, a BLASTn search revealed that none of the linear replicons contained a terminal Palindrome 1 sequence. By mapping Oxford Nanopore Sequencing (ONT) and Illumina reads to these replicons, we observed that the full archetypal telomere was often present in the sequencing data (Figure 3a), despite being absent in the assembly. Notably, the 13nt of the archetypal Palindrome I was only captured in the Illumina data, but not in the ONT data. We also noticed that some assemblies were truncated by hundreds to thousands of bases (Figure 3b) and would thus require further extension with ONT data to recover enough of the telomere for the terminal Illumina reads to map.

**Figure 3:**
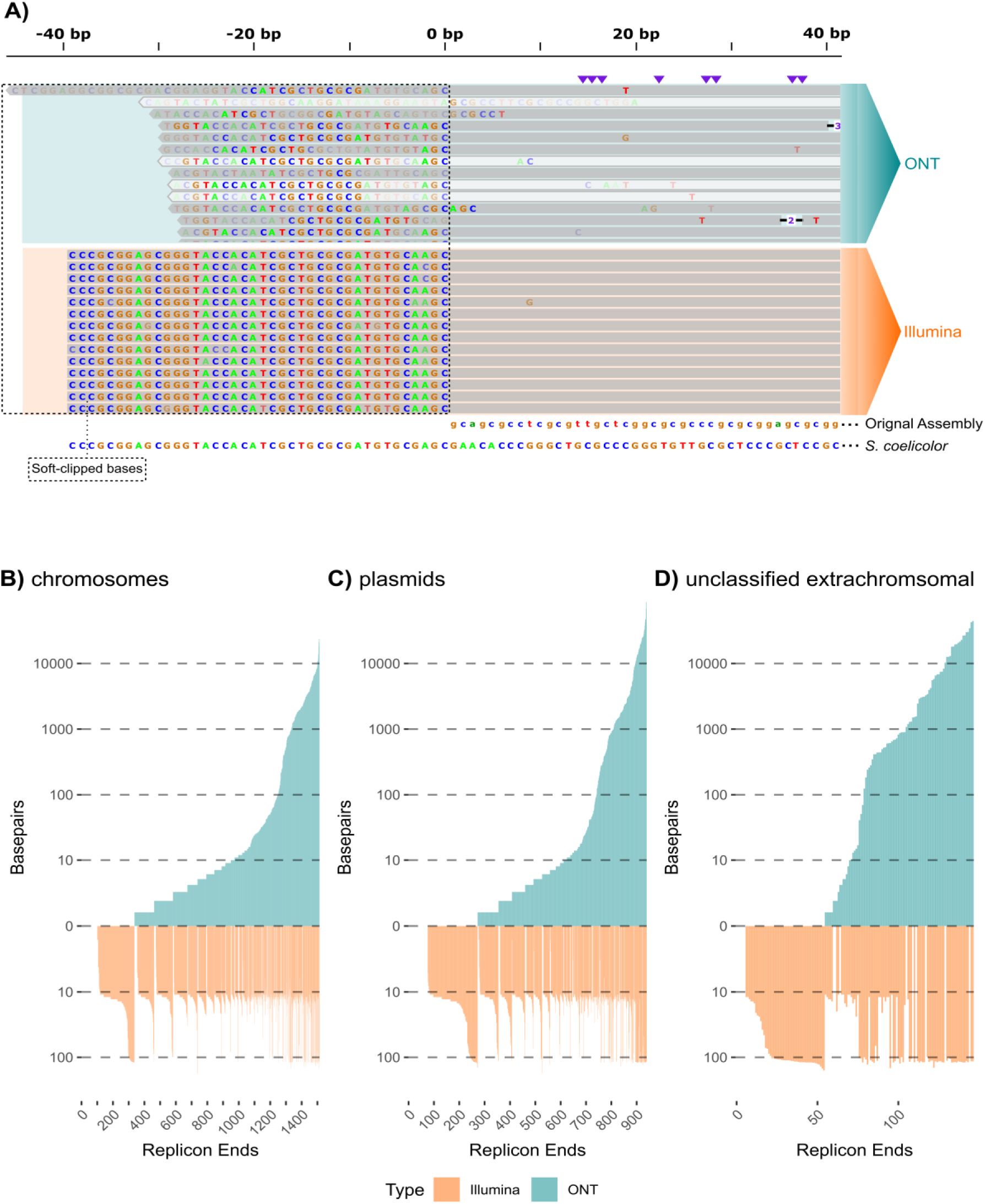
The terminal Palindrome I is missing in Oxford Nanopore data. **(A)** Oxford Nanopore Technologies (ONT) and Illumina reads mapped to the 5’-terminus of *Streptomyces niveus* NBC 01542 (accession: CP109389). ONT data is marked by a cyan background, while Illumina reads are marked by an orange background. Soft-clipped bases are highlighted with dotted lines. The original assembly sequence is shown in the bottom right corner, while the sequence of the *S. coelicolor A(3)* telomere, which carries an archetypal telomere, is shown below the assembly. (**B-D**) The result of extending linear G1034 replicons ends using Telomore, split across replicon types. Extensions from ONT are shown in cyan, while extensions using Illumina data are shown in orange. Replicons are sorted by the extension length using ONT. Each replicon occupies a single position, so ONT and Illumina extensions are shown together.

To automatically reconstitute the observed missing telomeric sequence we developed Telomore, which extends a linear replicon using either ONT or Illumina reads. Briefly, Telomore is a Python package that extends linear contigs using either terminally mapped ONT or Illumina reads (see method section for detailed description).

We used Telomore V0.3 (commit: #df04cfd) on 2601 linear replicons from the G1034 collection using first ONT and the Illumina reads to extend the assemblies. The logic behind this strategy is that ONT reads can typically extend a replicon end to within 10-15 nt of the terminus, after which Illumina reads can be used to complete these missing bases. After extension, we polished the assembly with Illumina reads to avoid the lower per base quality of ONT reads negatively affecting the quality of extended bases. The output from Telomore extension of linear replicon ends are depicted in Fig 3 B-D and Table 1 and 2.

Telomore extended 74% of replicon ends beyond the original assembly using ONT data. The median extension length of chromosomes and plasmids was only 4 nt and 5 nt, while the mean extension length was around 500nt and 1500nt. This is in line with our previous observation, where some outlier assemblies are missing hundreds of bases and thus require ONT to be fully extended, as the missing sequence exceeds the Illumina read length. Specifically, 20% of replicon ends were extended with >100 nt, while a few replicon ends were extended with up to 10 kb.

After ONT extension, Telomore extended 85% of replicon ends beyond the assembly with Illumina reads. Notably, the median extension length for chromosomes and plasmids were 12 nt and 13 nt, which closely resembles the sequence we observed to be missing in ONT data (Figure 3A). Since the Illumina reads used were 150 nt long, the maximal expected extension length is around 120 nt, allowing the final 30 nt to map to the assembly. For this reason, we expect replicon ends extended with >100 nt to potentially still be incomplete.

Unclassified extrachromosomal elements were in general extended more than plasmids and chromosomes. 52% of the unclassified extrachromosomal replicon ends were extended by >100 nt by ONT, while chromosomes and plasmids were only extended by >100 nt for 17% and 20% of replicon ends. Similarly, 48% of unclassified extrachromosomal replicon ends were extended with >100 nt using Illumina, while just 6% and 10% of chromosomes and plasmids are extended with >100 nt using Illumina. We suspect that this reflects that the unclassified extrachromosomal elements are more prone to contain misassemblies or be unresolved. Specifically, these are elements which could not be confidently classified as either chromosomes or plasmids in the original paper (9). This could be due to novelty, but could also be due to incompleteness.

After completing the sequences with Telomore, we repeated the BLASTn search for the archetypal Palindrome I and were able to detect it in 37% of the 2601 replicon ends. This demonstrates the power of Telomore to reconstitute telomeric sequence. Based on our observations, we suspected that replicon ends that had not been extended with Illumina data would be incomplete. Palindrome I was detected in 0% of the 184 ends extended only with ONT and 43% of the 2234 replicon ends extended with Illumina data. This indicates that while extension with ONT can help recover part of the telomere Illumina data is necessary to extend a replicon with the outermost bases.

#### Clustering of *Streptomycetaceae* telomeres

After reconstitution of the G1034 telomeres, we were interested in investigating telomeric diversity across *Streptomycetaceae and* assessing whether truncated telomeres in assemblies are a widespread problem in public genomes. It is, however, not common practice to submit raw sequencing reads along with a genome assembly. Therefore, we decided to collect all “complete” and “chromosome”-level *Streptomycetaceae* assemblies from. An overview of the number of replicon ends used in the clustering is available in Table 4 and a full overview can be found in supplementary Table S1. This allowed us to group the telomeres (including experimentally unknown classes), and to exclude incomplete or erroneous sequences.

**Table 4:** Overview of clustering of Refseq, Telomore-completed G1034, and known telomeres. The number of replicon ends in NCBI and Telomore completed G1034 is comparable, but the percentage which clusters differ markedly. The percent of replicon ends from Telomore completed G1034 which cluster, is similar to that of the known telomeres, whereas NCBI ‘complete’ and ‘chromosome’ level genomes rarely cluster. As a clustering control, circular sequences were included and as expected extremely rarely clustered with other sequences, many of which could be explained by wrongly assigned circularity. *notice that some sequences from the known telomeres column are also present in the NCBI complete RefSeq column as some chromosomes and plasmids have experimentally documented telomeres and are in RefSeq.

| **n=5324 total**<br>**seqs in uclust**<br>**file, of these**<br>**n=4606 linear** | **NCBI**<br>**complete**<br>**Refseq** | **Telomere**<br>**completed**<br>**G1034** | **Known**<br>**telomers \*** | **NCBI**<br>**complete**<br>**RefSeq** | **Telomere**<br>**completed**<br>**G1034** |
| --- | --- | --- | --- | --- | --- |
| topology | linear | linear | linear | circular | circular |
| total chromosomes | 651 | 759 | 83 | 27 | 3 |
| total plasmids | 313 | 473 | 42 | 113 | 188 |
| unplaced/extrachromosomal | 5 | 74 | 2 | 0 | 32 |
| total replicons | 969 | 1306 | NA | 140 | 223 |
| replicon ends | 1938 | 2601 | 127 | 280 | 446 |
| singletons | 64.3% (1246) | 15.1% (393) | 7.1% (9) | 82.1% (230) | 97.8% (436) |
| doubletons | 11.6% (225) | 6.9% (179) | 8.7% (11) | 11.1% (31) | 0.9% (4) |
| tripletons | 3.2% (62) | 1.3% (34) | 2.4% (3) | 1.1% (3) | 0% (0) |
| telogroups (4+ members of cluster) | 20.9% (405) | 76.7% (1995) | 81.9% (104) | 5.7% (16) | 1.3% (6) |

The sequence which harbors palindrome I-IV in the archetypal *S. coelicolor* telomeric system is crucial for telomere function and is 89 nt long (3), so we decided to use a sequence length of 90 nt for clustering and a 72% identity threshold. The identity threshold was decided after observing multiple instances of differences on both sides of palindrome stem sequences which would not affect the secondary structure of similar telomeres. After manual inspection of alignments and predicted loop structure we decided to discard clusters with three or fewer members.

### Most “complete” NCBI RefSeq assemblies do not contain two telomeres

Of the 4606 replicon ends, 54% (2467 replicon ends) were grouped into 137 clusters with four or more members, with the single largest telocluster containing 327 replicon ends. These clusters included 81.9% of known telomeres and 76.7% of the G1034 replicon ends. However, only 20.9% of replicon ends of complete NCBI Refseq assemblies fell within these clusters. A complete chromosomal sequence should be flanked by a telomere on each side. However, only 15% of NCBI Refseq chromosomes have two identified telomeres, compared to 78% of chromosomes in the Telomore-completed G1034 collection (Figure 4). These findings suggest that most NCBI Refseq assemblies are incorrectly annotated as complete, as they have truncated telomeres or unresolved terminal inverted repeats (TIRs). Since the TIR is often particularly difficult for assemblers to resolve (9), we speculate that many assemblies hosted at NCBI do not have fully resolved TIRs, which would also make it impossible to resolve both telomeres.

**Figure 4:**
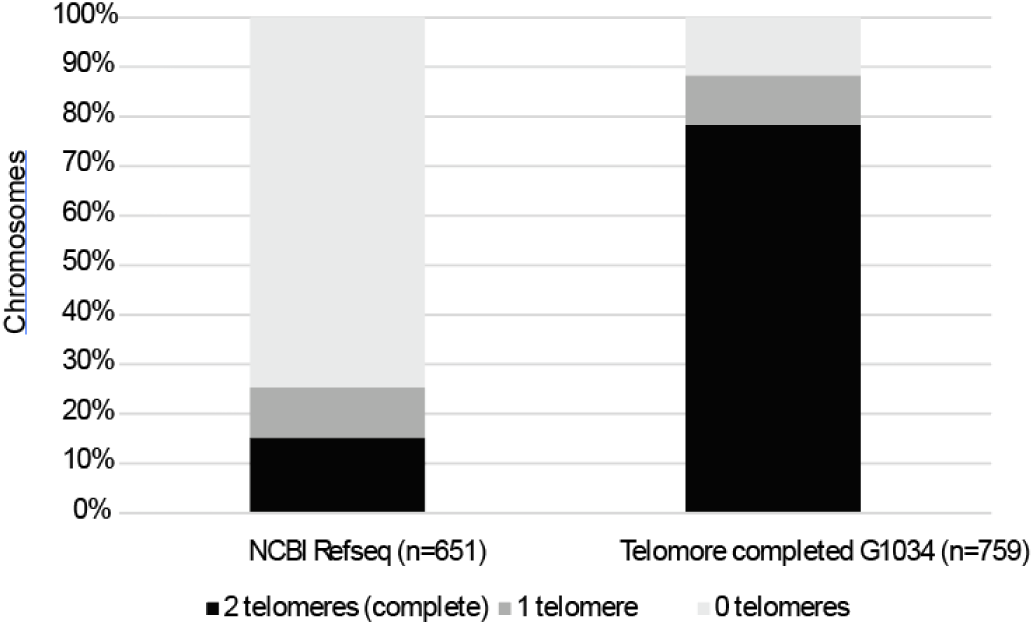
The chromosomes of most complete Refseq genomes does have two identifiable telomeres. Comparison of complete NCBI Refseq chromosomes with Telomore-complete G1034 by the number of chromosomes with zero, one or two telomeres assigned to a telocluster.

A disadvantage of clustering by similarity is that clustering in and of itself does not prove telomeric function, just similarity. As such, sequences that do not cluster could simply be particularly rare telomeres. However, we observe that the majority (74.1 %) of the discarded G1034 replicon ends are those that we expect to be truncated (see SI1: Clustering of circular elements). As such, we are confident that the majority of discarded sequences are truncated rather than particularly rare.

### The majority of identified clusters are archetypal

Out of the 137 telomere clusters, 62 clusters representing 56.2% of all clustered sequences contained the archetypal Palindrome 1. This is in line with previous findings showing that the archetypal telomere-type is the most widespread in streptomycetes (7). At the moment it is unclear why this telomere type is the most prevalent. Recently, telomeric transposons have been described in *Streptomyces* which are able to mobilize the telomere and facilitate telomere replacement(27). Certain clades of these transposons co-locate with *Tap-tpg*. Thus, frequent telomere replacement with the archetypal telomere might be one of the drivers making this the most prevalent type.

#### Telomeres of Kitasatospora and Embleya

The majority of available complete linear *Streptomycetaceae* genomes are from the genus *Streptomyces*. To explore the telomeres in non*-streptomyces* we investigated the telomere clusters identified for *Kitasatospora* and *Embleya*.

In *Kitasatospora*, which is closely related to *Streptomyces*, 78 replicon ends from 46 replicons are contained in nine clusters, some exclusively with other *Kitasatospora* sequences (33 replicon ends in cluster165, cluster194, and cluster328) and others where *Streptomyces* and *Kitasatospora* sequences clustered together (45 replicon ends in cluster4, cluster49, cluster50, cluster124, cluster245, and cluster303), in what is possibly the equivalent of the Sg2247 telomere class previously described. While all telomere clusters with *Kitasatospora* members have 3 or more palindromes, only one of them (cluster194, *Kitasatopora* replicon ends) contains the archetypal palindrome I, which could indicate that most *Kitasatospora* replicons have a different maintenance machinery than most *Streptomyces* replicons.

*Embleya*, with only two complete genomes currently available, has an unusual genome organization featuring a chromosome and an additional “secondary” chromosome (28).

Only 4 replicon ends from 2 plasmids were successfully extended with Illumina reads, both of which are singletons, which do not resemble a known class. While Telomore did not extend the chromosomes of *Embleya sp.* NBC_00888 *and Embleya sp.* NBC_00896, manual inspection of mapped nanopore and Illumina reads revealed that *Embleya sp.* NBC_0088 carries a telomere with a GC-repeat similar to Sg2247, while *Embleya sp.* NBC_00896 carries an archetypal telomere. This is in line with a preprint describing the evidence for a secondary chromosome in *Embleya australiensis* (28). We manually inspected the assembly from this strain and found that it contains an archetypal telomere.

##### The telomere maintenance machinery of Streptomycetaceae

After clustering the telomeres, we wished to investigate how the proteins that maintain the telomeres relate to these clusters. Currently, three pairs of telomere maintenance proteins have been characterized; Tap/Tpg, Tac/Tpc and GtpA/GtpB, which are all linked to particular classes of telomeres. There are, however, several classes for which the maintenance proteins are undescribed. Thus, we returned to the linear G1034 replicons and mined them for known telomere maintenance proteins using BLASTp before searching for potentially novel proteins in the strains where no known system was found.

We detected the presence of at least one full set of known telomere maintenance proteins in 76% of the 762 strains with linear genomes in the collection, or 52% of replicons. We expect that one system can be sufficient to maintain several replicons, as this has been previously demonstrated (29). As shown in Fig 5A, most strains encode homologs of Tap/Tpg, while few strains encode homologs of Tac/Tpc or GtpA/GtpB. All three systems are preferentially located within the first or last 10% of the replicon (Figure 5B). This preference is not observed for plasmids, which might reflect the smaller size of the plasmids compared to the chromosomes.

**Figure 5:**
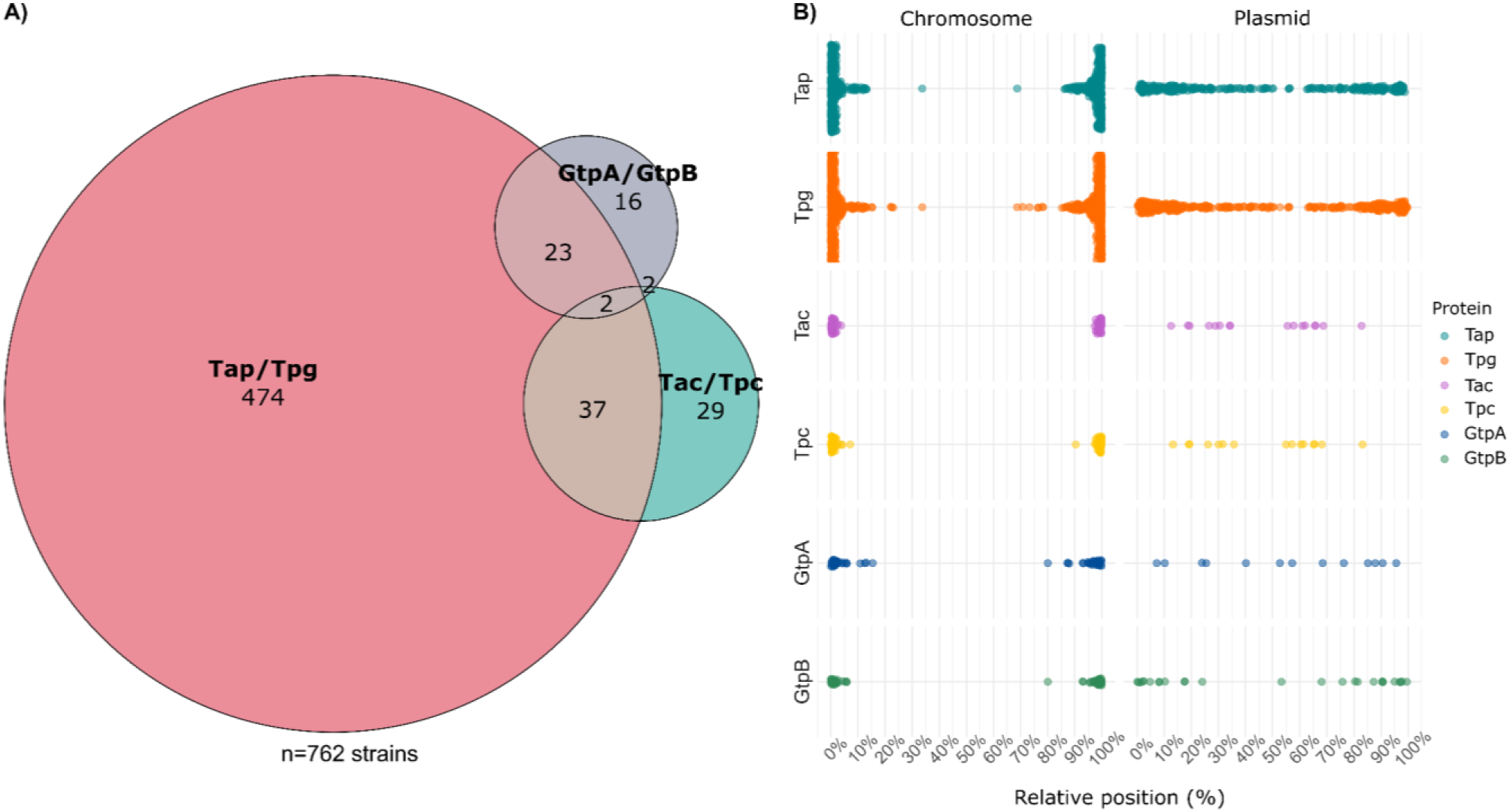
Know telomere maintenance proteins in the G1034 genomes. (**A**) Number of genomes with at least one full telomere maintenance system detected B) The relative genomic position of the genes encoding telomere maintenance proteins across all replicons. Unclassified extrachromosomal hits are not depicted as there were only 10 Tpg and 4 Tap hits.

The telomere maintenance systems are thought to be encoded as operons (1). Consistent with this we find that the median intergenic distance between the gene-pairs are 11 bases, 1 base and 1 base, for Tap/Tpg, Tac/Tpc and GtpA/GtpB respectively). In some cases, however, the distance is much longer and the pair not co-located and unlikely to be transcribed as an operon.

To investigate whether any particular maintenance system is associated with any particular telomeres, we clustered co-located proteins into “families” at 70% similarity and investigated their relationship to the 137 teloclusters described above.

### The non-archetypal systems

Tac-Tpc was found with teloclusters 11 (n=37), 26 (n=3), 111(n=10), 303 (n=4) and 328 (n=5), which all feature an initial 4-6 Gs and 4-nt hairpin loops as previously described in the literature for the SCP1 telomere class(30). GtpA-GtpB were found together with telocluster 2 (n=4), 11 (n=3) and 100 (n=21) where both telocluster 2 and 11 are linked to either Tap-Tpg or Tac-Tpc by many replicons not carrying GtpA-GptB, thus these clusters are likely not maintained by GtpA-GptB. Telocluster 100 features an initial 5’-CCG-3’ as well as loops of 5’-TCC-3’ or 5’-TCCC-3’ as previously described in literature for SG13350 (31).

### The archetypal system, Tap-Tpg

A total of 611 co-located Tap-Tpg gene-pairs were identified using the BLAST-based approach. Of these 89% (n=544) were found with teloclusters that contain the archetypal palindrome I, while 11% (n=67) were found with teloclusters that did not contain Palindrome I. 40 of these ends are found with teloclusters that co-occur more frequently with other systems and exhibit characteristics of them. Thus, these ends are not maintained by the Tap-Tpg pair and have at some point likely undergone telomere replacement. The remaining 27 are found with teloclusters that contain Palindrome I but have been extended additionally (telocluster 50 (n=24) and 644 (n=3)).

While the association between Tap and Tpg family is strong, such that a given Tap-family tend to co-locate with one or few specific Tpg-families, the relationship between Tap or Tpg family to telomere clusters is only moderate (Cramer’s V of 0.5). This is driven by the fact that most combinations of Tap and Tpg-families occur with several different telomere clusters. Most archetypal telomeres feature Palindrome II loops of 5’-TGC-3’ and Palindrome III loops of 5’-TGC-3’ or 5’-TCC-3’. Thus, it might not be surprising that they do not exhibit a strong specificity between most of the clusters.

Some archetypal clusters, however, do feature unusual Palindromes II and III. For example, telocluster 13 and 1870, which feature 5’-AGA-3’ (See Figure 6) loops in both Palindromes II and III. Telocluster 13 is exclusively linked to Tap-family 1 and Tpg-family 3 (n=10 G1034 replicons, n=17 G1034 replicon ends) and Tap-family 1 and Tpg-family 3 are exclusively linked to telocluster 13. Further, the AGA carrying telomere is plasmid specific both in G1034 genomes and RefSeq genomes (n=14 replicons and n=23 replicon ends). This telocluster includes pSCL3 (CM001018), one of the linear megaplasmids harboured by *Streptomyces claviligerus* ATCC 27064. The plasmids are diverse and range in size from 30 kb to over 400 kb, united mostly by conserved terminal repeats. In the larger megaplasmids, however, the conserved region extended beyond the telomere to incorporate a region of 50 kB ([Figure S1]). This region predominantly encodes for proteins involved in DNA maintenance and replication and the bacterial stress response, including homologues of LexA, TerD and DnaJ. With these findings in mind we speculate that this represents a distinct subset of the archetypal family of telomeres, similar to the system identified by Tidjani et al. (32). The identification of the AGA teloclusters represents the first description of a potentially plasmid specific *Streptomycetaceae* telomere, where the claim is supported by multiple independent observations and compared to chromosomal telomeres, which together makes a random genomic location unlikely.

**Figure 6:**
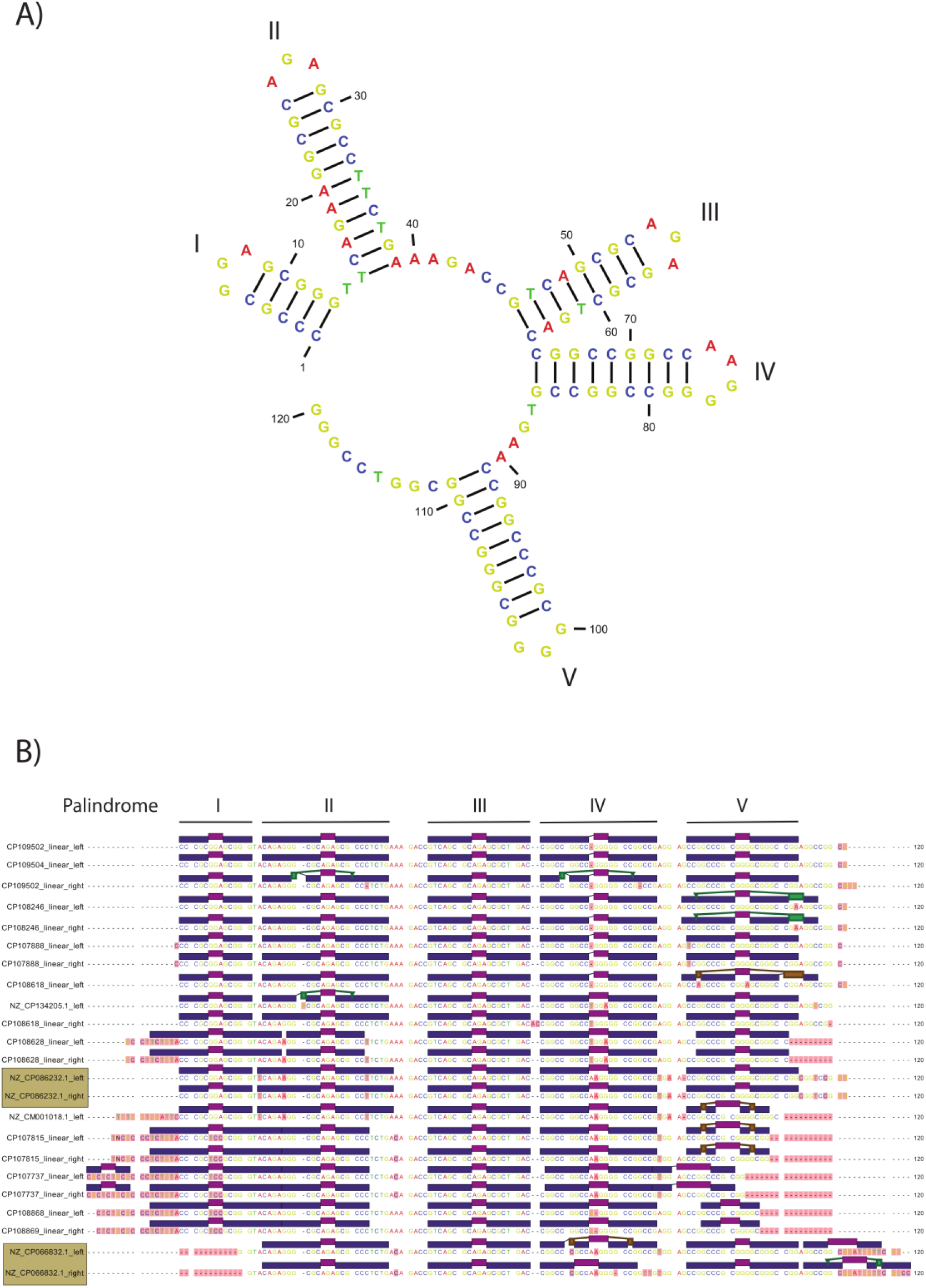
Discovery of a plasmid-specific archetypal telomere. **(A)** The predicted folding of *Streptomyces clavuligerus* strain DSM 738 plasmid pSCL3, featuring the 5’-AGA-3’ loop in the second and third palindrome. **(B)** An alignment of all members of telocluster 13 and 1870 (sand-coloured boxes). The annotation above each sequence is its predicted folding. Differences from the consensus sequence are highlighted in red.

### The first candidate for the maintenance of Sg2247-telomeres

We then wanted to identify potential candidates for the telomere maintenance systems of the 23% (n=179) of strains with no identified maintenance system. To achieve this, we collected the proteins from all the G1034 linear genomes, clustered them at 70 % identity, and sorted them into whether they had an identified telomere maintenance system (identified_system) or not (no_system). From this, we identified one protein family, which occurred in about half of the strains with no system identified and very few of those with an identified system (see Figure 7a).

**Figure 7:**
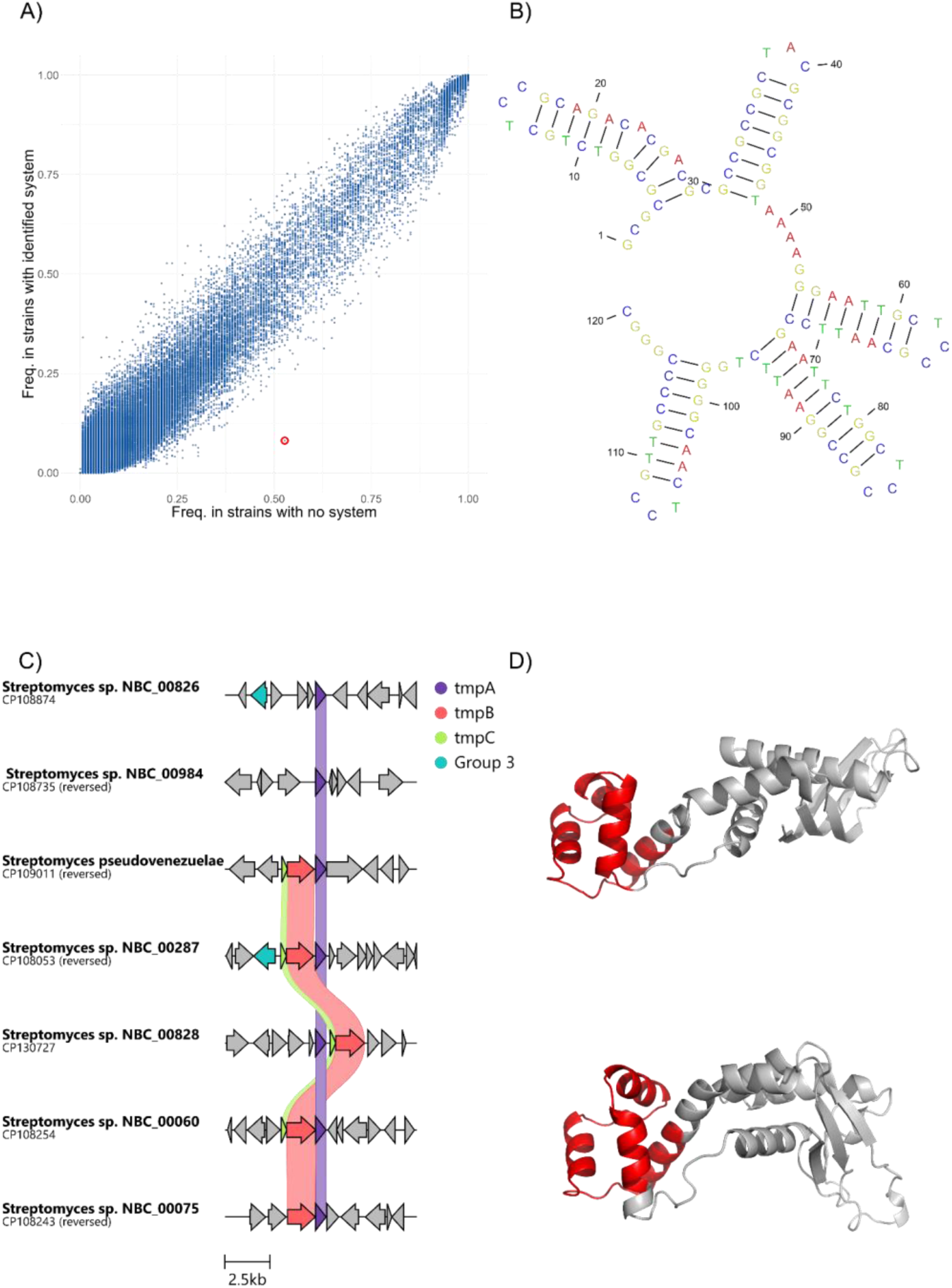
A candidate protein for maintenance of Sg2247 telomeres. **(A)** The frequency of protein families in the group with no identified system vs those without. The group represented by OHA19_44885 is marked with a red circle. **(B)** The predicted structure of the left-side of CP107889, a member of telocluster 18 showcasing a folding similar to that of Sg2247. **(C)** Examples of tmpA, tmpB and tmpC co-occurring with each other. **(D)** Top: The predicted fold of Tpg. Bottom: The predicted fold of TmpA. The red sections are highlighted based on their matching positions to the Lederbergvirus P22 C2-repressor (PDB-entry: 2r1j, D-chain) as determined by phyre2.

Members of this family, represented by OHA19_44885 from *Streptomyces sp.* NBC_00012, are preferentially encoded at the chromosomal arms and co-occur with teloclusters (12 (n=7), 18 (n=24), 40 (n=6), 67 (n=5), 99 (n=3), 161 (n=23), 224 (n=9), 240 (n=4), 253 (n=8), 326 (n=3), 342 (n=5), 729 (n=4), 748 (n=4)) that have the features of the Sg2247-class. Namely, an initial GCGCGC and loops of 5’-TCC-3’, 5’-TAC-3’ and 5-’TCC-3’. Thus, this protein family (hence forth tmpA (telomere maintenance protein A) may be responsible for maintaining telomeres of the Sg2247-class.

Previously characterized telomere maintenance proteins work in pairs. By inspecting the 5 kb up and downstream of *tmpA* genes, we observed that they were sometimes colocated with a larger helix-turn-helix domain protein (hence forth TmpB). This protein, like Tap, features an N-terminal Cro/C1 HTH domain. Searching the previously identified hits, showed that in 56% of the 125 hits, the two proteins were co-located. Thus, we hypothesize that the maintenance mechanism is similar to that of Tap-Tpg and Tac-Tpc, where one protein is used as a protein primer and deoxynucleotidylated by the other protein. In 36% of the identified hits, the proteins are also co-located with a smaller HTH-domain containing protein (tmpC). A similar pattern has been observed for a sub-type of the archetypal telomere by Tidjani et al (32), where Tap seems to be split into two proteins.

Although InterProScan did not identify any domains, TmpA had several high-confidence matches in the NCBI nr-proteins database for transcriptional regulator – a common misannotation for DNA-binding proteins. Tpg and TmpA were modelled using ColabFold version 1.55 (25) and aligned using TM-align, resulting in a score of 0.58 indicating that the two proteins adopt a similar fold. Visualizing the structure in Pymol shows that both proteins feature a helix-turn-helix domain (see Figure 7C). Taken together these findings support the role of the protein in DNA-binding.

## CONCLUSIONS

By analyzing the replicon ends of 762 linear genomes, we showed that ONT data does not capture the telomeres of linear *Streptomycetaceae* genomes, which lead to incomplete assemblies. We develop the tool Telomore to extend these incomplete genomes. We clustered the G1034 replicon ends with publicly deposited genomes and previously validated telomeres, to create a catalogue of >2000 telomeres in 137 similarity-based clusters. Using these clusters, we find that only 15% of NCBI RefSeq chromosomes have two identified telomeres indicating that most public genomes are incomplete. In comparison, 78% of the G1034 chromosomes extended by Telomore had two identified telomeres, showing that the strategy outlined can help improve completeness. By mining the G1034 genomes for maintenance proteins, we identify a known maintenance system in 76% of all strains. Mining the 24% of strains with no identified system, we identify a candidate protein family of DNA-binding proteins that we hypothesize is responsible for maintenance of Sg2245-class telomeres. Finally, we discover the first likely plasmid specific telomere type.

## ACKNOWLEDGEMENTS

We would like to thank Troels Østergaard Hansen for valuable discussion of assembly errors in *Streptomyces*.

## AUTHOR CONTRIBUTIONS

**David Faurdal**: Conceptualization, formal analysis, visualization, writing – original draft.

**Thom J. Booth**: Formal analysis, visualization, writing – review & editing.

**Tilmann Weber**: Supervision, funding acquisition, writing – review & editing.

**Tue Sparholdt Jørgensen**: Conceptualization, formal analysis, visualization, writing – original draft.

## SUPPLEMENTARY DATA CONFLICT OF INTEREST

None.

## DATA AND SOFTWARE AVAILABILITY

All raw Oxford and Illumina reads used in this study can be found under BioProject PRJNA747871. Scripts used to analyze data and produce figures can be found at https://github.com/dalofa/telomore_paper. Telomore is available as a python package/command-line tool at https://github.com/dalofa/telomore. The similarity-based clusters can be found in github repo: https://github.com/dalofa/telomore_paper.

