## Supporting Information for "Tying up loose ends: Recovering thousands of missing telomeres from Streptomyces and other Streptomycetaceae genomes"

**AUHTORS**

David Faurdal[1], T.J. Booth[1], Tilmann Weber[1*] and Tue Sparholt Jørgensen[1*]

[1]The Novo Nordisk Foundation Center for Biosustainability, Technical University of Denmark, Søltofts Plads, Building 220, 2800 Kgs. Lyngby, Denmark

#### Contents

SI 1: Clustering of circular elements ........................................................................... 2

Figure S1: Dotplots of telocluster 13 ........................................................................... 3

### SI 1: Clustering of circular elements

Because clustering of replicon ends only imply sequence similarity but not telomeric function, we wanted to explore to which degree circular sequences would cluster. Because circular plasmid contigs are expected to break at a random point, and not harbor telomeres, and rarely be rotated to a certain gene, our expectation was that they would not cluster with telomeres. For both NCBI and G1034 sequences, the large majority of artificial circular replicon 'ends' aka contig start and ends, did not cluster, with 81.9% and 97.8% singletons, respectively, and further 12% and 1% respectively in clusters with two or three members. Interestingly, when diving into the 6 circular G1034 replicon ends which did cluster in groups with 4 or more members, we found that the topology assigned did not match the assembly graph topologies, and that they all in fact should be assigned linear topology based on this. Similarly, the very prominent Streptomyces collinus Tu 365 plasmid pSCO1, where the telomere has been analyzed several times, was also assigned circular topology, despite it being linear. These errors have now been fixed.

A caveat to the clustering strategy is that circular replicons are often rotated to dnaA for chromosomes, and replication genes for plasmids, which could skew the analysis, and indeed, of the 21 NCBI circular contig ends clustering in groups of four or more, 14 can be confidently classified as telomeric (meaning that the topology is wrongly assigned in RefSeq), and six of the remaining seven circular contig ends are dnaA related (dnaA for lefthand side, upstream dnaA for righthand side). Interestingly, in nine cases these dnaA sequences cluster with supposedly linear streptomyces chromosomes, which seem to have large scale assembly artefacts. We speculate that these nine chromosomes (NZ_CP046623.1, NZ_CP030862.1, NZ_CP047146.1, NZ_CP136652.1, NZ_CP169412.1, NZ_CP016279.1, NZ_CP029241.1, NZ_CP134875.1, NZ_CP016279.1), though linear, were incorrectly 'rotated' to dnaA in an automated workflow. All dnaA related sequences and their mate in the other contig end were discarded before further analysis.

### Figure S1: Dotplots of telocluster 13

Figure S1: Dotplots showing conserved regions between *Streptomyces clavuligerus* ATCC 27064 plasmid pSCL3 (CM001018.1) and select other members of telocluster 13: a) *Streptomyces* sp. BB1-1-1 plasmid unnamed2 (CP134205.1), b) *Streptomyces* sp. NBC 00829 plasmid unnamed1 (CP108868.1), c) *Streptomyces xanthophaeus* strain NBC 01137 (CP108628.1), and d) *Streptomyces avidinii* strain NBC 01157 plasmid unnamed2 (CP108618.1). The locations of the Tap-Tpg homologues are highlighted in sky blue.

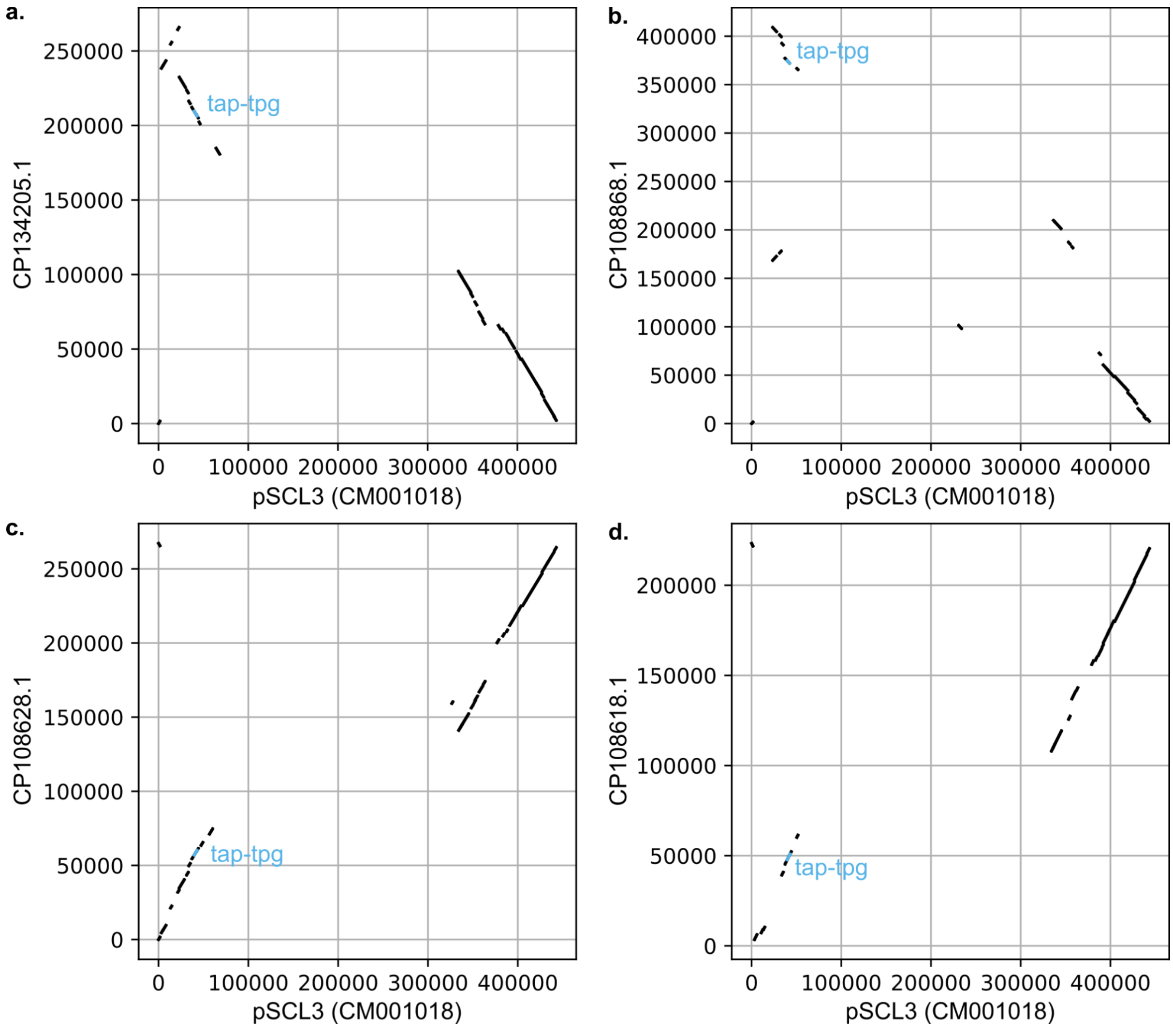
